# Label-free characterization of Amyloid-β-plaques and associated lipids in brain tissues using stimulated Raman scattering microscopy

**DOI:** 10.1101/789248

**Authors:** Volker Schweikhard, Andrea Baral, Vishnu Krishnamachari, William C. Hay, Martin Fuhrmann

**Affiliations:** Leica Microsystems CMS GmbH, Am Friedensplatz 3, 68165 Mannheim, Germany; Neuroimmunology and Imaging Group, German Center for Neurodegenerative Diseases (DZNE), Bonn, 53127, Germany

## Abstract

The brains of patients with neurodegenerative diseases such as Alzheimer’s Disease (AD) often exhibit pathological alterations that involve abnormal aggregations of proteins and lipids. Here, we demonstrate that high-resolution, label-free, chemically-specific imaging using Stimulated Raman Scattering Microscopy (SRS) provides novel insights into the biophysical properties and biochemical composition of such pathological structures. In brain slices of a mouse model of AD, SRS reveals large numbers of Amyloid-β plaques that commonly form a characteristic, three-dimensional core-shell structure, with a fibrillar proteinaceous core surrounded by a halo-like shell of lipid-rich deposits. SRS spectroscopic imaging allows for a clean, label-free visualization of the misfolded (β-sheet) Amyloid-β content in the plaque core. Surrounding lipid-rich deposits are found to contain comparatively high concentrations of membrane lipids (sphingomyelin, phosphatidylcholine), but lower levels of cholesterol than healthy white matter structures. Overall, the SRS spectra of plaque-associated lipids closely resemble those of nearby neurites, with the notable difference of a higher degree of lipid unsaturation compared to healthy brain structures. We hypothesize that plaque-associated lipid deposits may result from neuritic dystrophy associated with AD, and that the observed increased levels of unsaturation could help identify the kinds of pathological alterations taking place. Taken together, our results highlight the potential of Stimulated Raman Scattering microscopy to contribute to a deeper understanding of neurodegenerative diseases.

## Introduction

Alzheimer’s Disease (AD) is a chronic neurodegenerative disease that is characterized by neurological symptoms such as progressive cognitive impairment and altered behavior, accompanied by neuronal cell death and the buildup of senile plaques and other pathological changes in certain areas of the brain. And yet, despite decades of intensive research, the molecular and cell biological mechanisms of disease progression remain incompletely understood, and there is currently no efficient treatment to cure, halt, or slow down the progression of the disease. An important hallmark of AD pathology is the aggregation of misfolded Amyloid-β (Aβ) peptides into large abnormal *extracellular* deposits. Furthermore, it is commonly believed that neurodegeneration in AD is linked to the presence of small Aβ oligomers in the *intracellular* space that interfere with neurotransmission. However, the aberrant formation, transport, and defective clearance of these neurotoxic Aβ species in brain tissues remains incompletely understood. Several hypotheses suggest that plaque-associated lipids may act as surfactants that mediate an exchange of Aβ between aggregated forms that are sequestered in the extracellular plaques and soluble small oligomeric forms that lead to toxicity. However, the origins and roles of these plaque-associated lipids in the progression of the disease remain poorly understood. As a consequence, there is a high demand for new analytical methods that can probe the interplay between brain lipids and Aβ with high spatial resolution and chemical specificity in intact brain tissues.

Coherent Raman Scattering (CRS) microscopy denotes a set of nonlinear optical microscopy techniques, most prominently Coherent Anti-Stokes Raman Scattering (CARS) and Stimulated Raman Scattering (SRS), that image biological structures by exploiting the characteristic, vibrational contrast of the sample molecules^1,2,3^. The optical setup and principle of SRS microscopy are illustrated in **Supplemental Figure S1** and described in Materials and Methods. In recent years, CRS has emerged as a powerful approach for label-free, biochemical imaging of neuronal structures and for dynamic monitoring of functional parameters such as membrane potential^4^ and neurotransmitter concentrations^5^. In addition, CRS has been used in fundamental research on neurodegenerative diseases, to characterize Alzheimer’s pathology^6,7,8^, Parkinson’s Disease^9^ and multiple sclerosis^10^, or to study peripheral nerve degeneration in ALS^11^. Currently, there is a major push to develop label-free histopathological analyses based on CRS (sometimes integrated in a multi-modal approach with other nonlinear optical imaging techniques such as two-photon fluorescence or harmonic-generation microscopy)^12^, with major efforts in brain cancer^13,14,15^, other cancers^16,17,18^ and a range of other diseases^19,20^. Beyond disease-diagnosis, SRS has been applied in fundamental research on disease *mechanisms*, e.g. to link aberrant cholesterol metabolism pathways to prostate cancer aggressiveness^16^ and to identify lipid desaturation as a marker for cancer stem cells^17^.

Here, we demonstrate the potential of SRS to provide label-free, high-resolution biochemical and biophysical characterizations of healthy and diseased brain tissues. Our results provide novel insights into Amyloid-β plaque biology and highlight the potential of coherent Raman scattering microscopy to contribute to a deeper understanding of neurodegenerative diseases.

## Results

### Label-free SRS microscopy of diseased brain slices reveals lipid-rich pathological deposits associated with Aβ plaques

We applied SRS microscopy to unstained brain slices from mice carrying an amyloid precursor protein and a presenilin 1 mutation that exhibit AD-like pathology^21,22^, as well as from healthy control mice. We first acquired large-area overview tile-scan SRS images of healthy and diseased brain slices (**Figure 1 (A)** and **(B)**, respectively). In these images, SRS contrast arises from CH_2_ stretch vibrations at 2850 cm^-1^ predominantly found in lipids. Lipid-rich white matter regions (bright horizontal bands at image bottom) are easily discernible, as well as the less lipid-rich grey matter of the cortex (darker green regions in the top half of the images). In diseased brain slices, additional, localized lipid-rich pathological structures are seen, which are absent in the healthy cortex. Further detailed analysis described below demonstrates that these localized pathological structures correspond to lipid deposits associated with Amyloid-β plaques. **Figure 1 (C)** shows a high-resolution two-color SRS image of such an Amyloid-β plaque (red, SRS at 1678 cm^-1^ as explained below), and lipid structures (green, SRS at 2850 cm^-1^). Plaques are found to exhibit a characteristic, three-dimensional core-shell structure, in which a fibrillar core is surrounded by a halo-like arrangement of lipid-rich structures. Orthogonal slice views through a 3D image stack, and a movie of the stack, are shown in **Supplemental Figure S2** and **Supplemental Movie S1**, respectively. In the healthy tissue regions surrounding the plaque, SRS reveals myelinated neurites (bright tubular structures) and neuronal cell bodies (dark regions devoid of lipids) embedded in homogenous grey matter. The observation of prominent plaque-associated lipid deposits raises questions: What is the source of these lipids, and how do these structures form? What is their role in AD progression? In the following, we present a detailed characterization of Aβ plaques and associated lipids, utilizing the quantitative spectroscopic imaging capabilities of SRS, to begin to address these questions.

**Figure 1:**
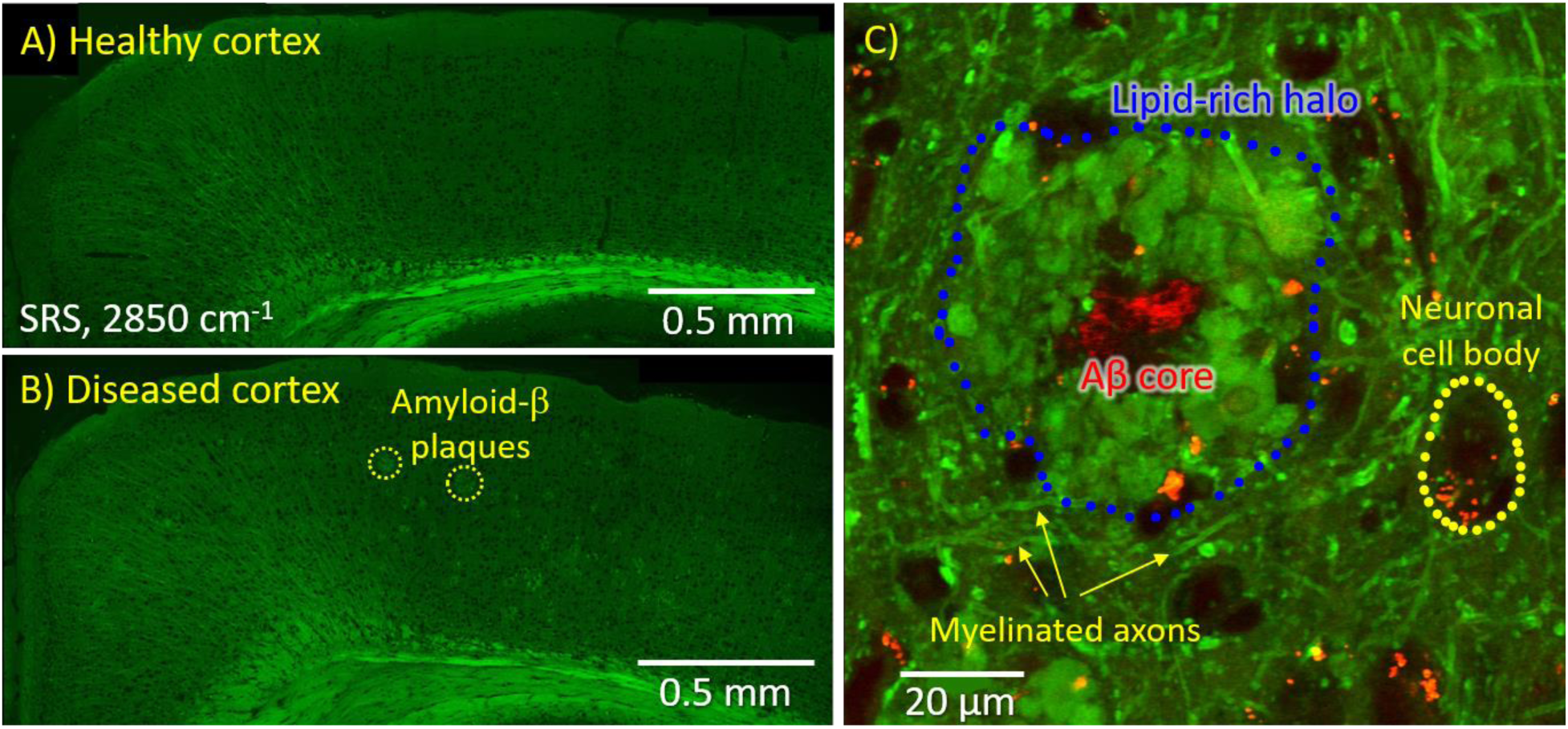
(A, B) Label-free, large-area overview images of unlabeled mouse brain slices, acquired using Stimulated Raman Scattering microscopy (SRS). Signals arising from CH2 stretch vibrations at 2850 cm^-1^ predominantly found in lipids are shown in a healthy (A) and a diseased (B) mouse cortex. (C) High-resolution, two-color SRS image of an Amyloid-β plaque (red, SRS at 1678 cm^-1^), and associated lipid deposits (green, SRS at 2850 cm^-1^).

### High-resolution hyperspectral SRS microscopy of healthy and diseased brain structures

To characterize the biochemical composition of Amyloid-β plaques, we performed hyperspectral SRS microscopy, in which SRS images are acquired sequentially over a large range of wavenumbers. As a result, a full SRS spectrum with a spectral resolution of ∼12 cm^-1^ is available for each image pixel. Importantly, it has been shown that the resulting SRS spectra are fully equivalent to spontaneous Raman spectra and therefore allow for similar image analysis and quantification procedures developed for spontaneous Raman data. CARS spectra, by contrast, suffer from spectral distortions and off-resonant background signals that make a quantification considerably more difficult.

Representative example images in **Figure 2 (A, C)** show a tissue region at the interface between the cortex (bottom of image) and white matter (top). Representative spectra of white matter, grey matter, cell nuclei, Aβ plaque cores and plaque-associated lipid halo regions are shown in **Figure 2 (B)**. Note, the spectral data shown in **Figure 2** are raw data that did not require any post-processing (*i*.*e*., no background subtraction and no smoothing), and were acquired on regular fused silica coverslips. This is a major advantage of coherent Raman imaging, in which the signal is generated selectively in the laser focus, over spontaneous Raman imaging, which suffers from large spectral backgrounds originating from substrates and/or sample mounting compounds, and hence require extensive data subtraction procedures and the use of specialized, expensive substrates that produce lower background signals (MgF_2_, CaF_2_).

**Figure 2:**
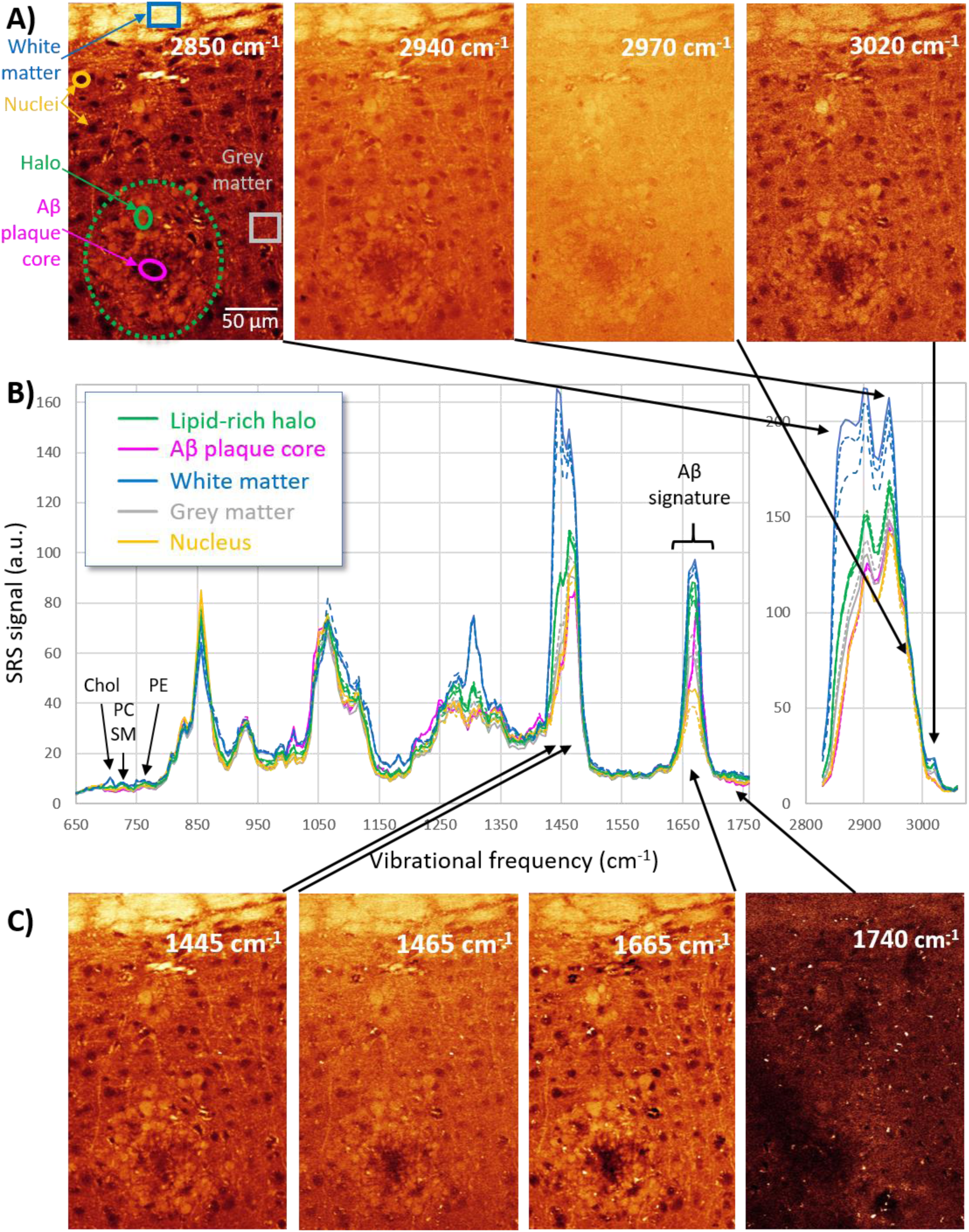
Hyperspectral SRS microscopy applied to a diseased mouse cortex. SRS images are acquired sequentially at a range of wavenumbers, as in the representative images shown in (A,C). As a result, a full SRS spectrum is available for each image pixel. Representative SRS spectra of the brain structures indicated in (A) are shown in (B). 2-3 spectra from different instances of the same structure are shown to indicate typical spectral variation. For detailed explanations and analysis of the “Aβ-signature”, see **Figure 3**. For “Chol, PC/SM, PE”, see **Figure 4**.

SRS spectra comprise two regions, (i) the high-wavenumber region (^∼^ 2800-3100 cm^-1^) with signals from CH_2_ and CH_3_ stretch vibrations representing the sample’s total protein and lipid contents (2940, 2850 cm^-1^, respectively), and =C-H vibrations associated with unsaturated lipids (∼3020 cm^-1^); and (ii) the fingerprint region (∼500-1800 cm^-1^) containing large numbers of peaks that are often unique for a given compound.

Notable features in the fingerprint spectra of brain slices include (i) the near-absence of C=O stretch signal from oxidized lipids (esters) around 1740 cm^-1^, (ii) a prominent peak around 1660 cm^-1^ comprising the Amide I C=O vibration of proteins and C=C stretch vibrations in unsaturated lipids and sterols. (iii) a prominent double peak around 1465 cm^-1^ (Amide II in proteins) and 1445 cm^-1^ (total lipid signal from CH_2_ bending), (iv) 1200 - 1400 cm^-1^ region comprising the Amide III mode from proteins and further lipid signatures (1300, 1268 cm^-1^), (v) 1020-1150 cm^-1^ signals from carbohydrates, (vi) a sharp peak at 1005 cm^-1^ from Tryptophan, (vii) peaks around 850 and 900-950 cm^-1^ mainly from carbohydrates and nucleic acids, and (viii) a region from 600-800 cm^-1^ containing weak but distinct signals that allow the distinction of several major lipid classes (Cholesterol at 707 cm^-1^, membrane lipids – mainly phosphatidylcholine (PC) and sphingomyelin (SM) at 723 cm^-1^, as well as phosphatidylethanolamine (PE) around 760 cm^-1^). Note, SRS spectra are highly reproducible between different brain slices imaged on different days, as demonstrated in **Supplemental Figure S3**, and therefore even small variations in the spectra can contain highly relevant information. In the following, we examined the biochemical and biophysical information contained in several of these features in more detail.

### SRS allows for a clean, label-free visualization of Aβ-peptide aggregates localized in the plaque cores

We first sought to identify a way of directly and specifically visualizing the aggregated Aβ content of the pathological structures. We focused on the Amide I (C=O stretch) vibration, which was shown to be particularly sensitive to protein secondary structure in previous spontaneous Raman measurements^23,24^ and coherent Raman imaging studies^6,7^.

In **Figure 3**, we present a hyperspectral SRS analysis of the Amide I spectral region. Images taken at 1665 cm^-1^ show the overall tissue architecture, with contrast contributed both from the Amide I mode of proteins and from the overlapping C=C stretch vibration of unsaturated lipids. These images display a notable absence of signal in the central “core” of the pathological structure that is enclosed by the lipid deposits. When tuning the SRS contrast to 1675 cm^-1^, bright fibrillar structures appear in this location, whereas contrast in the remainder of the tissue is somewhat weakened. We assign these fibrillar structures and their blue-shifted Amide-I mode to Amyloid-β aggregates. This assignment agrees with a characteristic ∼10 cm^-1^ spectral blue-shift of the Amide I mode for proteins folded into a β-sheet conformation that was found in previous spontaneous Raman measurements on AD-affected tissues^23,24^. We validated the Amyloid-β specificity of the 1675 cm^-1^ peak by co-localization with fluorescent signals from Methoxy-04 (MX04), a molecule that binds to peptides in β-sheet conformation and is commonly used to stain Aβ plaques in Alzheimer’s research (see **Supplemental Figure S4**). Our assignment also agrees with results from two previous studies on AD-affected tissues^6,7^, where similar structures exhibiting the same frequency-shift were observed and validated as Aβ plaques using fluorescent dye labeling with thioflavin S, another stain commonly used to visualize Aβ.

**Figure 3:**
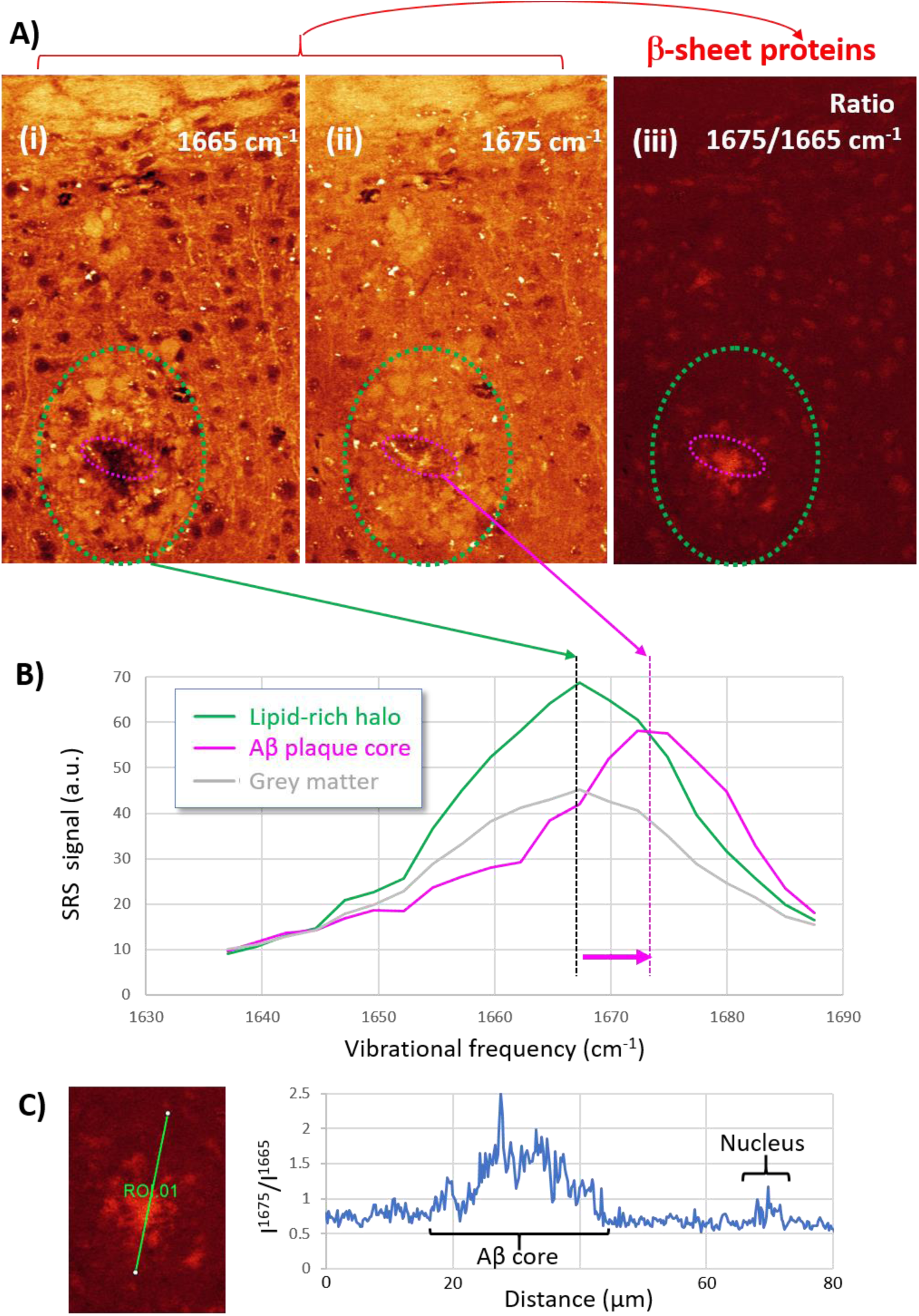
Hyperspectral SRS microscopy of the Amide I band reveals Aβ-aggregates in the plaque core. (A) (i) SRS images at 1665 cm^-1^ (Amide I vibration in regular proteins, as well as C=C stretch in unsaturated lipids) exhibit a dark void in the core of the Aβ plaque (pink line). (ii) At 1675 cm^-1^ (shifted Amide I vibration in β-sheet proteins), strong signals from fibrillar structures appear in the plaque core, consistent with the known misfolded β-sheet structure of Aβ-aggregates. (iii) Ratiometric visualization of Aβ plaques. The ratio of intensities at 1675 and 1665 cm^-1^ is shown, providing a clean visualization of the plaque core. (B) SRS spectra of healthy grey matter, lipid-rich regions in the plaque halo and core region are shown, demonstrating the specific frequency shift of the Amide I mode in the plaque core compared to regular proteins in the surrounding grey matter. (C) Line profile of the intensity ratio I^1675^/I^1665^ through the Aβ core and a nearby cell nucleus.

The ratio of images at taken at 1675 cm^-1^ and 1665 cm^-1^, shown in **Figure 3 (A)** (iii), therefore provides a clean and specific visualization of Aβ aggregates in plaques on an otherwise homogeneous background. We note that cell nuclei are visible with a very weak contrast in this image as well. This effect is seen in healthy brain slices too, and should therefore not be interpreted as evidence for the presence of Aβ in cell nuclei. **Figure 3 (C)** shows a line profile of the intensity ratio I^1675^/I^1665^ through the Aβ plaque core and a nearby cell nucleus, demonstrating the high specificity of this intensity ratio for the Aβ core.

### Aβ plaque associated lipid deposits are rich in membrane lipids, but low in cholesterol

The lipidome of the healthy brain is known to be composed largely of glycerophospholipids, sphingolipids, and cholesterol, which localize predominantly to neuronal membranes and myelin^25^. However, several contradicting hypotheses exist as to the origin of plaque-associated lipids in AD. We therefore decided to investigate the spectral range from ∼700 – 770 cm^-1^ in detail, which contains characteristic fingerprint signatures for some of the major lipid classes^26^ (**Figure 7**). The sterol ring breathing mode from cholesterol is found at 707 cm^-1^. While images at this wavenumber show that plaque halos (image area encircled in green) do contain some cholesterol, it is present there at substantially lower levels than in white matter regions at the top edge of the image. By contrast, plaques are comparatively rich in membrane lipids (sphingomyelin SM and phosphatidylcholine PC, 725 cm^-1^). A signature from phosphatidylethanolamine (PE) is seen in the spectra as well (feature around 745-775 cm^-1^). Consistent with published spectra^26^, the PE peak at 760 cm^-1^ is substantially broader than the ones of cholesterol and SM/PC. Images at this wavenumber suggest that PE is more diffusely distributed throughout the entire tissue region, with the highest PE concentrations found in white matter, and a notable exclusion of PE in the plaque core. Note, while the spectral signatures from cholesterol, PC, SM and PE in the 695-775 cm^-1^ region investigated here are very weak compared to the other signals examined in this work, all spectral features are highly reproducibly observed in very similar ratios in multiple different brain slices imaged on different days, as shown in **Supplemental Figure S5**.

Overall, plaque-associated lipids (green spectra) display an almost identical spectral signature to nearby neurites (yellow spectrum) in the spectral region shown in **Figure 4**, but differ significantly from the spectra of cholesterol-rich white matter (See also **Supplemental Figure S6** for the full spectral range). Taken together, our observations suggest that membrane components from nearby neurites, rather than cholesterol-rich white matter pools, appear to be the likely source of plaque-associated lipids.

**Figure 4:**
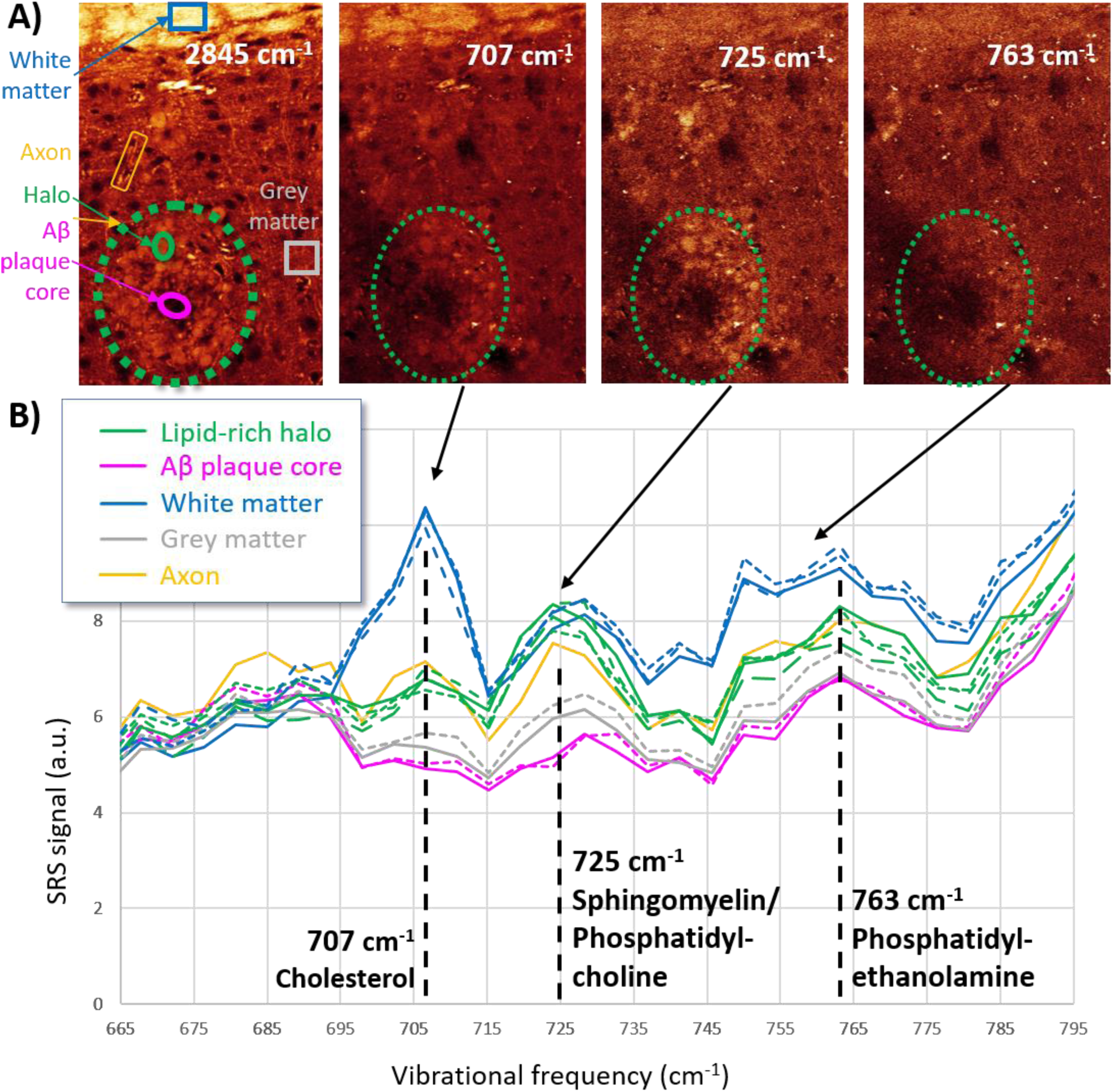
Stimulated Raman Scattering microscopy probes the composition of plaque-associated lipids and healthy brain structures. SRS images (A) and spectra (B) of the sterol ring breathing mode (707 cm^-1^), choline group vibrations (725 cm^-1^) and the ethanolamine group vibration from phosphatidylethanolamine (763 cm^-1^) are shown. An image of the total lipid content acquired at 2845 cm^-1^ is shown for reference in the first panel.

### Plaque-associated lipids exhibit higher degrees of unsaturation than nearby white matter

Two further pathologically relevant biochemical metrics accessible with SRS are the degrees of lipid unsaturation and oxidation in tissues. In pure lipid samples, the ratio of C=C stretch signal at ∼1665 cm^-1^ to CH_2_ bending signal at ∼1445 cm^-1^ has been widely accepted as a quantitative metric for the degree of lipid unsaturation. Three independent calibrations, presented by three different research groups^26,27,28^ are in agreement that the average number of C=C bonds per lipid molecule N_(C=C)_ is linearly related to the intensity ratio I^1665^/I^1445^ = 0.60*N_(C=C)_. In samples containing proteins in addition to lipids, some care has to be taken before attempting a ratiometric quantification of lipid unsaturation, since the 1665 cm^-1^ peak also contains signals from Amide I protein vibrations, and the 1445 cm^-1^ lipid peak overlaps in part with the Amide II peak from proteins at 1465 cm^-1^. **Figure 5 (A)** shows data from brain slices in this spectral region. In the brain, a relatively clean spatial separation into lipid-rich structures and protein-dominated grey-matter tissue provides a way of eliminating protein signals from SRS spectra, by subtracting the spectra of nearby grey-matter regions from those of more lipid-rich structures. The result of this subtraction is shown in **Figure 5 (B)** and **Supplemental Figure S6**, and represents spectra of the lipid component of these structures with relatively little remaining contamination from protein signals. Quantification of the degree of unsaturation in different brain structures then yields N_(C=C)_ = 1.55(19) in plaque halos (average and standard error of 23 halo regions), much higher than in nearby white matter where N_(C=C)_ = 0.66(6) (average over 3 large regions), and in nearby unaffected axons where N_(C=C)_ = 0.90(45) (average over 6 axons).

**Figure 5:**
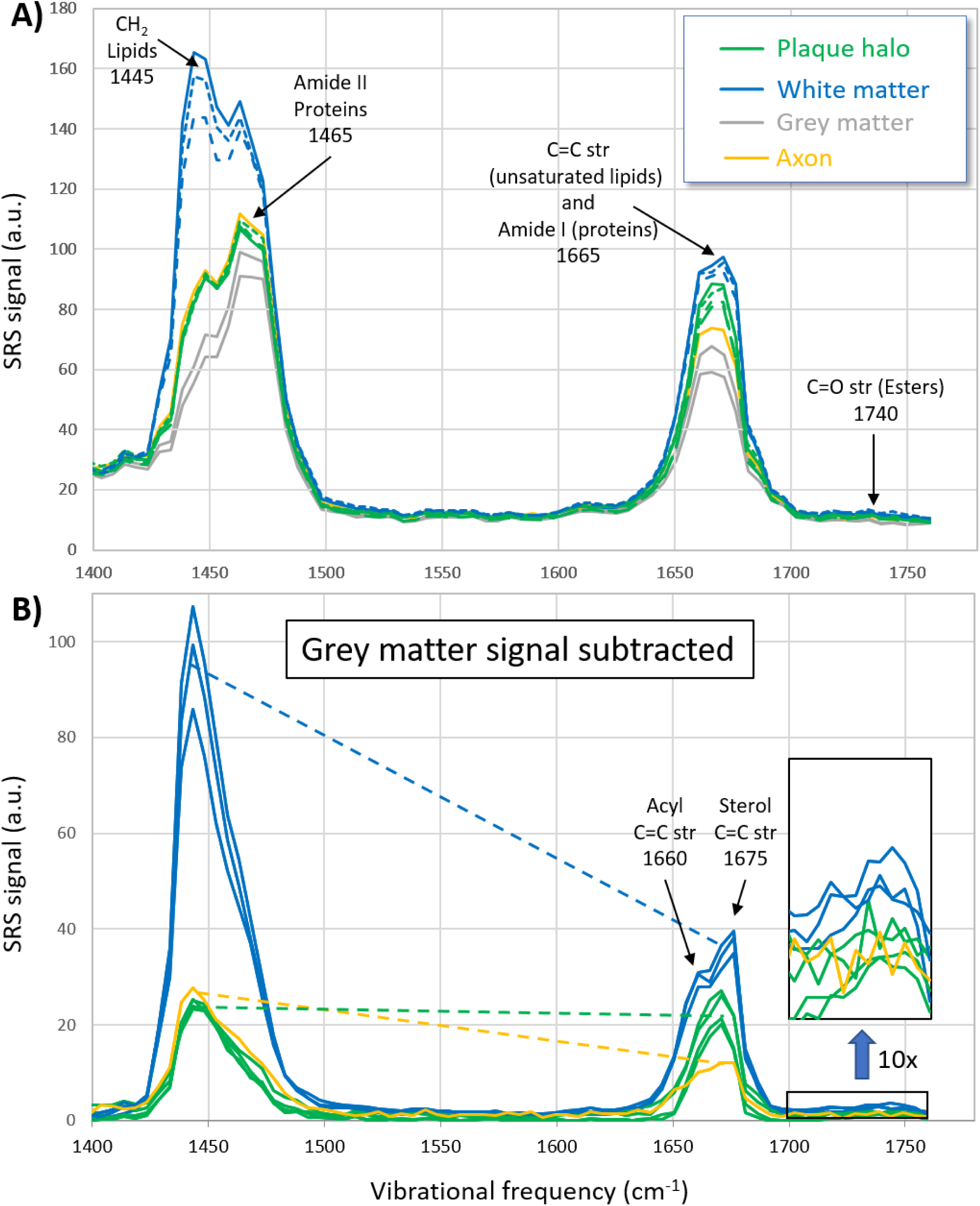
Ratiometric spectral analysis of lipid unsaturation. (A) Raw SRS spectra of the CH2 bend and C=C stretch bands. (B) “Lipid-only” SRS spectra of brain structures, obtained by subtracting the spectra of nearby grey matter from the full spectra of white matter, plaque lipids, and axons. The ratio of SRS intensities at 1665 and 1445 cm^-1^ (I^1665^/I^1445^) is directly proportional to the degree of lipid unsaturation. As indicated by the slope of the dashed lines, this ratio is higher in the plaque-halo compared to surrounding unaffected white matter and myelin-rich neurites.

Note, a ratiometric analysis of unsaturation is possible using high-wavenumber spectra as well, (see **Supplemental Figure S7**) and supports the finding of higher unsaturation levels in plaque lipids compared to healthy structures. The ratio of intensities at 3020 cm^-1^ to 2850 cm^-1^ from grey-matter-subtracted high-wavenumber spectra yields N_(C=C)_ ≈ 1.6 in plaque halos, N_(C=C)_ ≈ 0.5 in white matter, and N_(C=C)_ ≈ 0.6 in nearby axons. Note that signals from the C=C bond in the sterol ring of cholesterol contribute to the I^1665^/I^1445^ measure of lipisd unsaturation, whereas cholesterol does not have a strong signal around 3020 cm^-1^; hence, the qualitative agreement of high-wavenumber and fingerprint metrics for unsaturation rules out a dominant role of the sterol C=C vibration in our assessment of unsaturation.

An interesting feature of the “grey-matter-subtracted spectra” in Fig. 5 (B) is the splitting of the C=C peak around 1670cm^-1^ in white matter into a doublet (1660 and 1675 cm^-1^). This splitting is explained by a spectral shift between C=C vibrations in acyl and sterol groups^29^, both of which appear to be present in large concentrations in white matter. Note, the observed clear Cholesterol ring breathing signature in white matter at 707 cm^-1^ in **Figure 4** and the strong sterol C=C stretch in **Figure 5**, in combination with the near absence of 1740 cm^-1^ signal expected from cholesteryl esters, suggests that cholesterol is present in its unesterified form, consistent with prior observations in brain tissue^30^.

Interestingly, the near-absence of lipid C=O stretch signals at 1740 cm^-1^ of plaque-associated lipids also demonstrates that oxidation levels remain low even in these pathological structures. This observation is somewhat surprising, since oxidative stress is a key feature in the Alzheimer’s disease (AD) brain and has been found to manifest as lipid peroxidation^31^.

## Discussion

Label-free SRS imaging of brain slices from a transgenic mouse model of Alzheimer’s Disease reveals large numbers of Aβ aggregates, which we find are consistently surrounded by a halo of lipid-rich deposits. Similar associations of lipids with Aβ plaques have been observed in earlier studies using coherent Raman scattering^6,7^, imaging mass spectrometry^32^ and electron microscopy^33^, albeit with much lower spatial resolution or with a lack of chemical specificity. Interactions of lipids with pathological Aβ structures are under active investigation for their relevance to AD disease progression, because lipids are thought to have the capacity to extract small Aβ oligomers from aggregates in a detergent-like fashion^34^, thereby facilitating their transport to the intracellular space where they become neurotoxic. In addition, several major lipid classes are known to be disrupted in AD, including cholesterol, sphingolipids, phospholipids, and glycerolipids^25^. And yet, despite their obvious importance, the precise roles of pathological lipid alterations in the progression of AD have remained unclear.

Several hypotheses, not necessarily mutually exclusive, may explain the *origin* of plaque-associated lipids: (i) the *cholesterol hypothesis* states that cholesterol may play a specific role in Aβ plaque formation. This hypothesis is supported by the finding from mass spectrometric analyses that cholesterol and its transporter apoE are colocalized with Aβ plaques in transgenic mouse models of AD^35^. The mechanism by which cholesterol could be involved in AD pathology is however still unclear^25^. (ii) the *neuritic dystrophy hypothesis* proposes that plaque-associated lipids may originate from the degeneration of neurites and myelin sheaths in AD^36^. And, (iii) the *immune system hypothesis* suggests that plaque-associated lipids may originate from interactions of the brain’s innate immune system with pathological Aβ aggregates^37,38, 39^. In particular, plaques have been found to be tightly enveloped by microglia, which were found to assemble a barrier with a strong impact on plaque composition and toxicity^38^. Insights into the validity of either one of these hypotheses could open new avenues of investigation into the *roles* of the lipids in disease progression, and could potentially lead to the identification of new therapeutic targets for AD.

Our findings on the composition of pathological lipid structures are most in line with the neuritic dystrophy hypothesis: A comparison of the full SRS spectra of plaque-associated lipids to those of healthy brain structures reveals a high degree of similarity to the spectra of nearby neurites. In particular, a strong SRS signature at 723 cm^-1^ suggests the presence of high concentrations of membrane lipids (phosphatidylcholine and sphingomyelin). By contrast, relatively weak signals from the sterol ring breathing mode at 707 cm^-1^ show that plaque-associated lipids contain less cholesterol than surrounding healthy white matter structures, disfavoring a dominant role of cholesterol metabolism and transport in the formation of these structures. A notable feature in the spectra of plaque-associated lipids is the high degree of unsaturation compared to those found in healthy brain structures: both, a ratiometric analysis of signals at 1665 cm^-1^ (C=C stretch, unsaturated lipids) and 1445 cm^-1^ (CH_2_ twist, total lipids), and the strength of the 3020 cm^-1^ =C-H stretch signal from unsaturated lipids compared to the total lipid signal at 2850 cm^-1^ (CH_2_ stretch) suggest that plaque-associated lipids exhibit higher degrees of unsaturation than white matter and neuritic structures, which themselves are already known to be very rich in (poly)unsaturated fatty acids. We hypothesize that these unique spectral signatures could help identify the kinds of pathological alterations taking place in neuritic structures as a consequence of AD.

## Conclusions

We have applied a label-free chemical spectroscopic imaging approach based on Stimulated Raman Scattering microscopy to probe Amyloid-β plaques in brain tissue slices from a transgenic mouse model of Alzheimer’s Disease. Our results show that plaques consistently form a core-halo structure, with a dense proteinaceous Aβ core surrounded by a halo of lipid-rich deposits. SRS enables a clean, label-free differentiation of the Aβ core from the regular protein content of the surrounding unaffected brain tissue, by exploiting a spectral shift of the Amide I vibration that arises from the misfolding of aggregated Aβ into a β-sheet conformation. A spectroscopic analysis of the surrounding plaque-associated lipid deposits reveals that these contain high concentrations of the membrane lipids phosphatidylcholine and sphingomyelin. By contrast, the concentrations of cholesterol in plaques are low compared to those in nearby healthy lipid-rich brain structures. Overall, the spectroscopic similarity of plaque-associated lipids to nearby unaffected neurites favors the hypothesis that pathological lipid deposits form around the Aβ core as a consequence of neuritic dystrophy processes associated with AD.

In the future, we anticipate that the spectroscopic imaging capabilities of SRS will enable similar, hypothesis-generating and -testing research in many areas of the life sciences and medicine^40,41^, ranging from fundamental research in cell and tissue biology, to preclinical research on disease mechanisms, to applications in histopathology and diagnostic imaging, and the identification of novel biomarkers for a wide range of diseases. The enormous information content of SRS spectroscopic imaging also highlights the potential of this technique as a versatile platform for label-free, high-content screening applications.

## Materials and Methods

### Transgenic mouse model

Mice that exhibit AD-like pathology were prepared as previously described^21,22^: Wild-type mice were crossbred with APP/PS1dE9 mice (MMRRC strain 034832) that expressed the human Swedish mutation within the Aβ domain of the amyloid precursor protein (APPK594N/M595L). Additionally, these mice carried a transgene encoding human presenilin 1 with a deletion of the exon 9. Both transgenes were expressed under the prion protein promoter^21^.

### Brain slice preparation

As described in a previous publication^22^, mice were deeply anesthetized with an i.p. injection of ketamine/xylazine (0.26/0.02 mg/g) and transcardially perfused with PBS (pH 7.4) followed by perfusion with 4% paraformaldehyde (PFA) for five minutes each. Subsequently, the brains were removed and post fixed overnight in 4% PFA at 4°C. Next, 50-100 μm thick brain sections were cut on a vibratome (Leica VT1200S). Un-stained slices were mounted between standard microscope slides and coverslips for SRS and CARS imaging.

### SRS and CARS Microscopy

Coherent Raman Scattering Microscopy (CARS and SRS) was performed on a Leica SP8 CARS laser scanning microscope with SRS option (Leica Microsystems, Mannheim, Germany). **Supplemental Figure S1** shows a schematic of the beam routing. Briefly, CRS signals are excited using two temporally and spatially overlapped pulse trains from a PicoEmerald S optical parametric oscillator (APE, Berlin, Germany). The pump beam wavelength is fixed at 1031.25 nm, and the Stokes beam is tunable from 720 – 980 nm, allowing the excitation of vibrations in the range of 4200 to 500 cm^-1^. Both pulses are ∼2 picoseconds in duration, providing a ∼12 cm^-1^ spectral resolution of the total system. For SRS microscopy, the Stokes beam intensity is modulated at 20 MHz using an Electro-Optical Modulator (EOM). To acquire SRS signals, the Pump beam intensity is recorded in the forward direction using a silicon photodiode, and demodulated using a Lock-In amplifier (Zürich Instruments, Zürich, Switzerland). CARS microscopy does not require intensity modulation, and signals are detected on photomultiplier tubes in the transmitted and epi-detected directions. Second harmonic generation (SHG) / two-photon excited fluorescence signals can be recorded simultaneously with the CARS signals. Forward SRS and Epi-CARS/SHG detection are possible simultaneously. Visible-light confocal laser scanning microscopy was performed on the same instrument, and used e.g. for Methoxy-04 co-staining experiments.

