## Supporting Information for "Label-free characterization of Amyloid-β-plaques and associated lipids in brain tissues using stimulated Raman scattering microscopy"

### Supplemental Figure S1

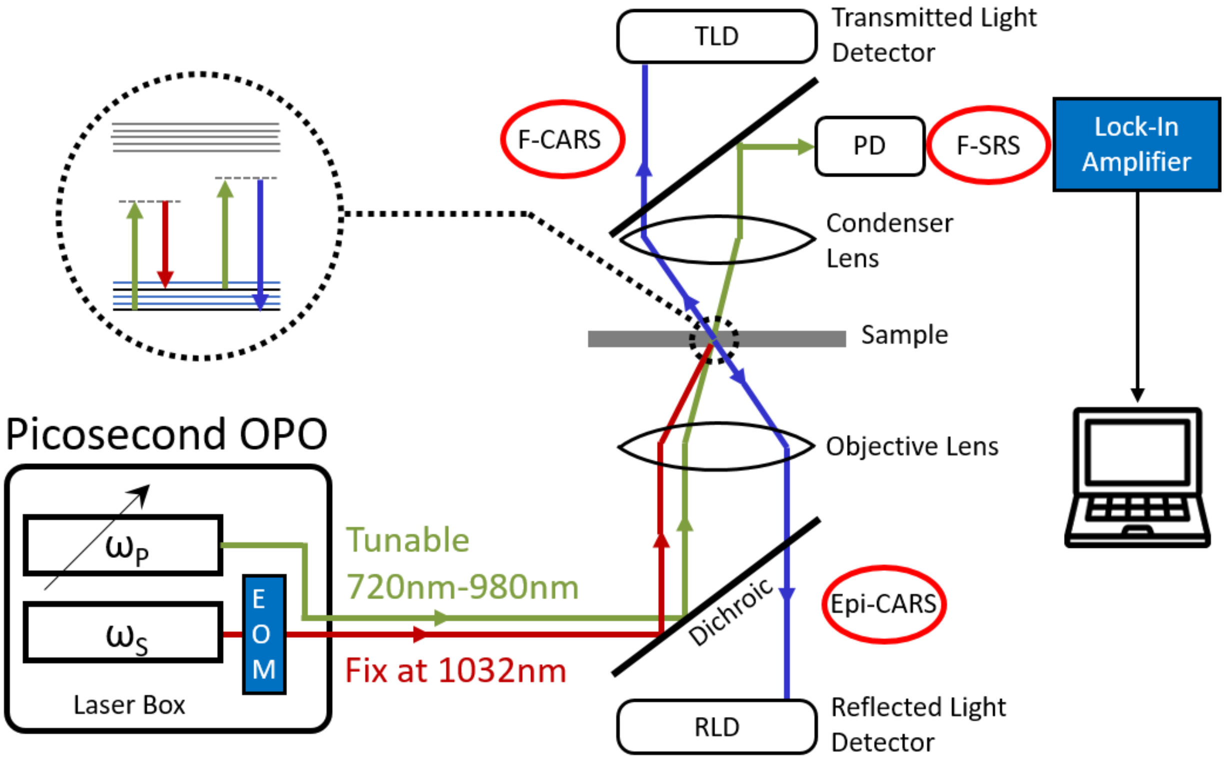


**Supplemental Figure S1:** Schematic of the beam routing used to excite CARS and SRS signals in the Leica TCS SP8 CARS laser scanning microscope platform with SRS add-on.

### Supplemental Figure S2

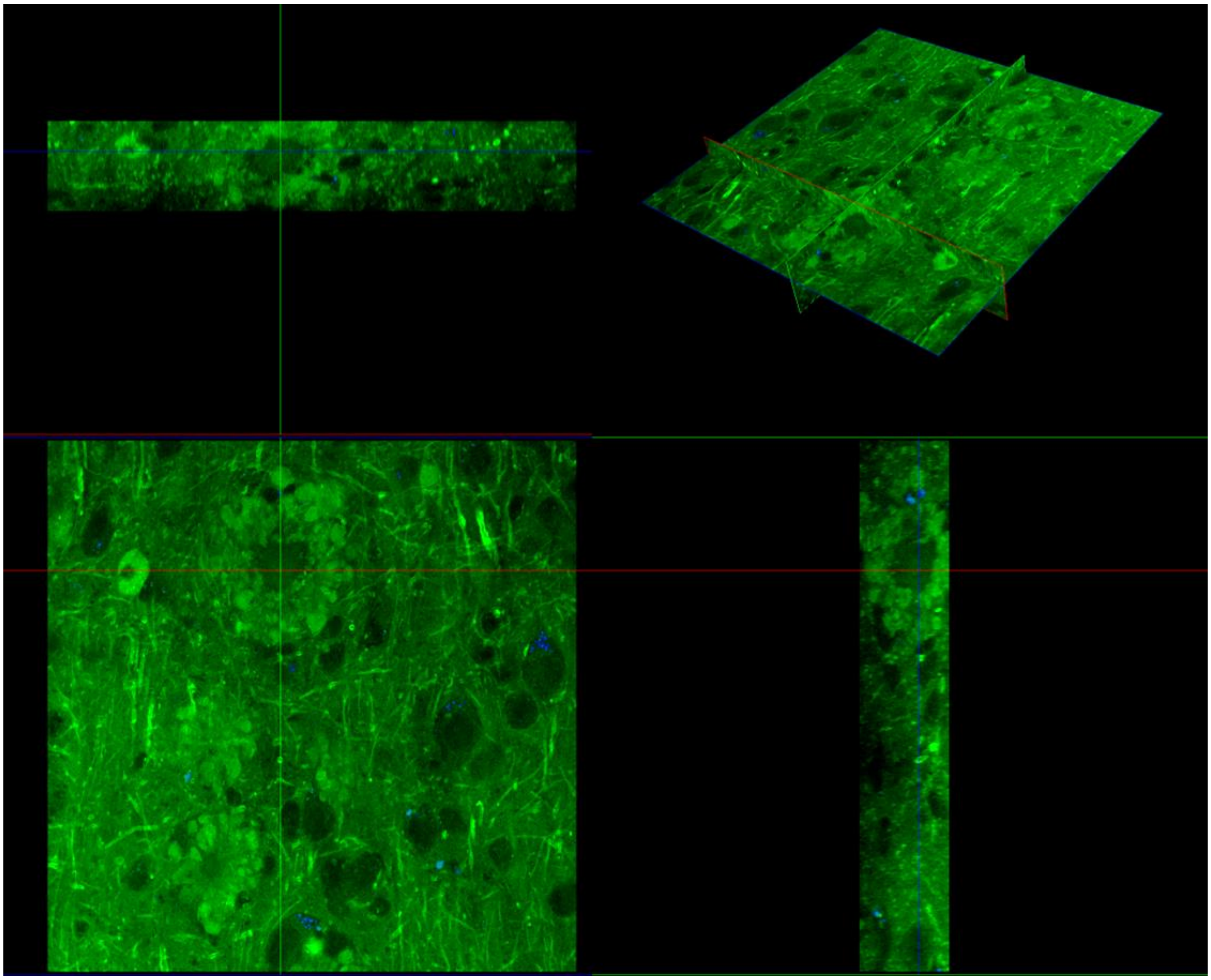

**Supplemental Figure S2:** Orthogonal slice views of a 3D image stack (200×200 µm xy image)×(33 µm z-stack) showing several Aβ plaques. The three-dimensional core-shell structure of plaques is visible. Green: Lipids (CARS at 2850 $cm^{-1}$); Blue: Two-photon excited autofluorescence.

### Supplemental Movie S1:

### See separate file “CARS2850 AF2850 Plaques 3D Movie.avi”

### Supplemental Figure S3

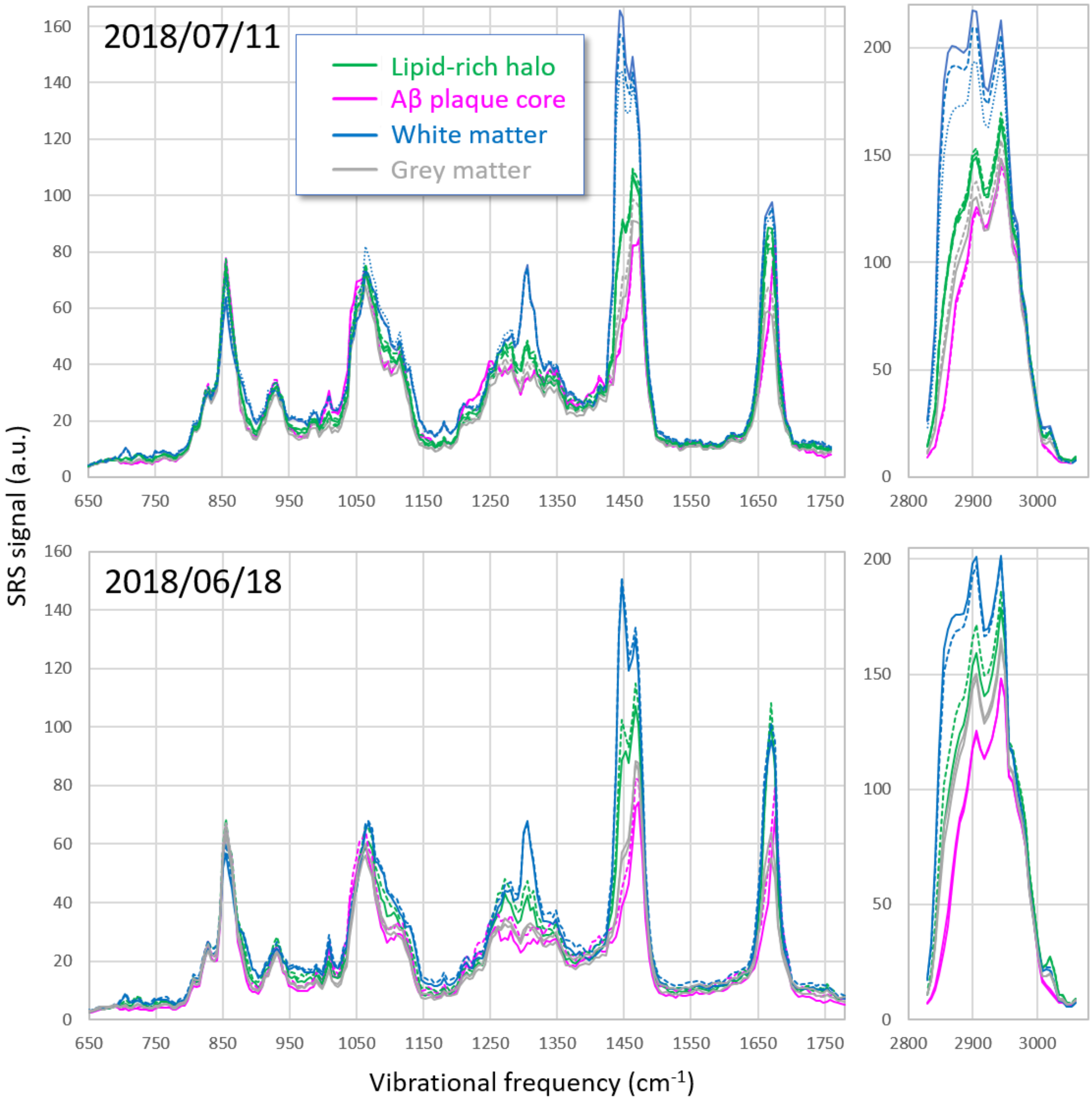


**Supplemental Figure S3:** Reproducibility of SRS spectra taken on different brain slices, on different days. Shown are raw data, without any background subtraction or smoothing.

### Supplemental Figure S4

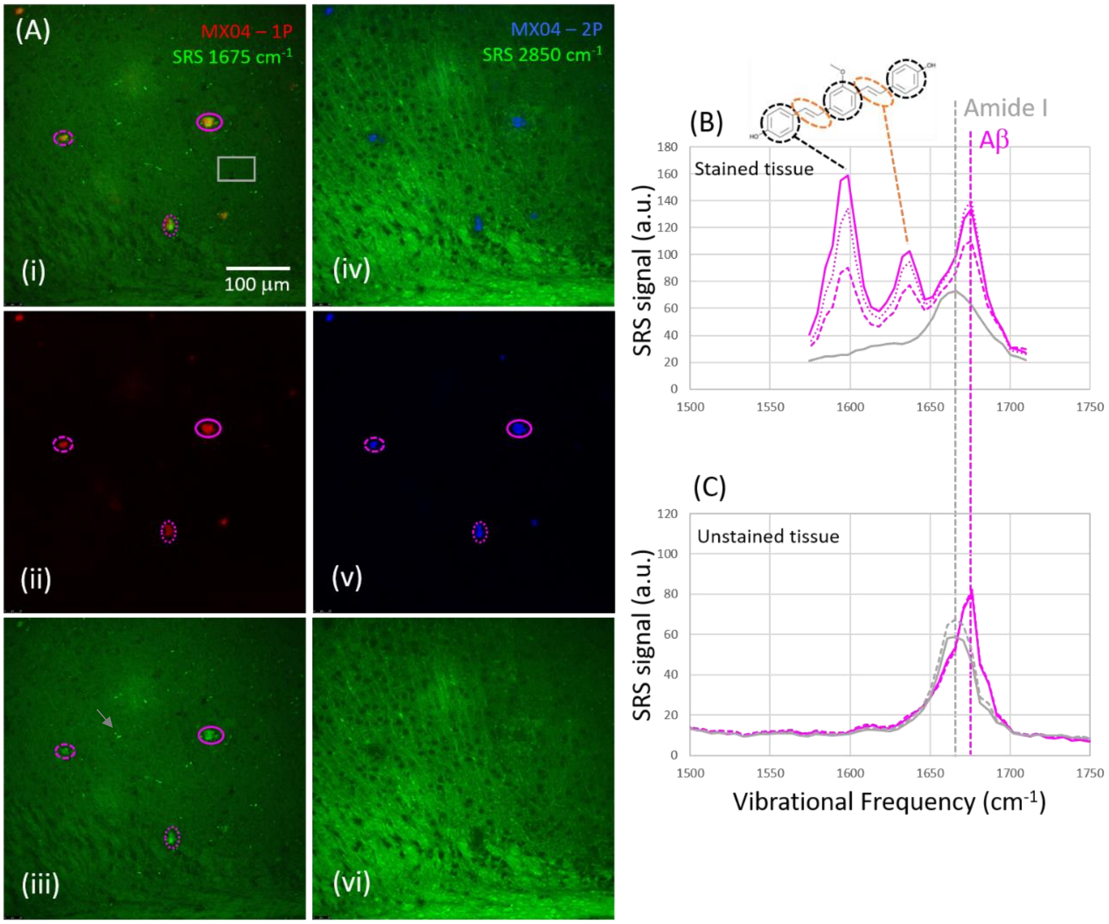


**Supplemental Figure S4:** Validation of the Amyloid-β specificity of the 1675 $cm^{-1}$ SRS resonance by co-localization with Methoxy-04 (MX04). **(A)** Overlays of SRS images and MX04 fluorescence: **(i-iii)** SRS images at 1675 $cm^{-1}$ and single-photon fluorescence of MX04 excited at 405 nm (red). The bright, localized structures in the 1675 $cm^{-1}$ SRS images (pink regions marked in **(iii)**) that we assign to Aβ-aggregates, perfectly colocalize with MX04 signals **(ii)**, confirming our assignment. Note, the additional smaller punctate structures visible in **(iii)** (e.g., grey arrow) do not exhibit any resonance in SRS spectra, and correspond to autofluorescent structures, not Aβ. **(iv-vi)** *Two-photon* fluorescence of MX04 (blue) can be excited by the Pump laser (~900 nm) and detected in the epi direction (380-550 nm detection window), and is shown here together with lipid signals (SRS 2850 $cm^{-1}$, green). Halo-like lipid structures **(vi)** are found surrounding the MX04 signals **(v)**, consistent with our observation that lipids surround the Aβ-plaque cores in unstained samples. **(B, C)** Detailed spectral analysis of SRS signals from MX04-stained **(B)** and unstained tissues **(C)**. The spectral features in the region between 1650-1700 $cm^{-1}$ show a close correspondence between stained and unstained tissues: The Amide I peak of regular proteins at 1665 $cm^{-1}$ (grey spectrum from the grey image region in **(A)**) and the frequency-shifted resonance of Aβ-aggregates at 1675 $cm^{-1}$ (pink spectra from pink regions in **(A)**). Below 1650 $cm^{-1}$, two new peaks appear selectively in the MX04-stained regions (pink) but not outside the stained regions (grey) or in unstained tissues. We assign these new peaks to vibrations of the MX04 molecule itself, i.e., the phenol ring at 1599 $cm^{-1}$ (https://www.chemicalbook.com/SpectrumEN_108-95-2_Raman.htm), and the C=C stretch of the ethylene group at 1637 $cm^{-1}$ (e.g., the C=C bond of the ethylene group in Trans-Stilbene has a similar chemical environment as in MX04, and has its resonance at 1641 $cm^{-1}$; https://www.chemicalbook.com/SpectrumEN_103-30-0_Raman.htm).

### Supplemental Figure S5

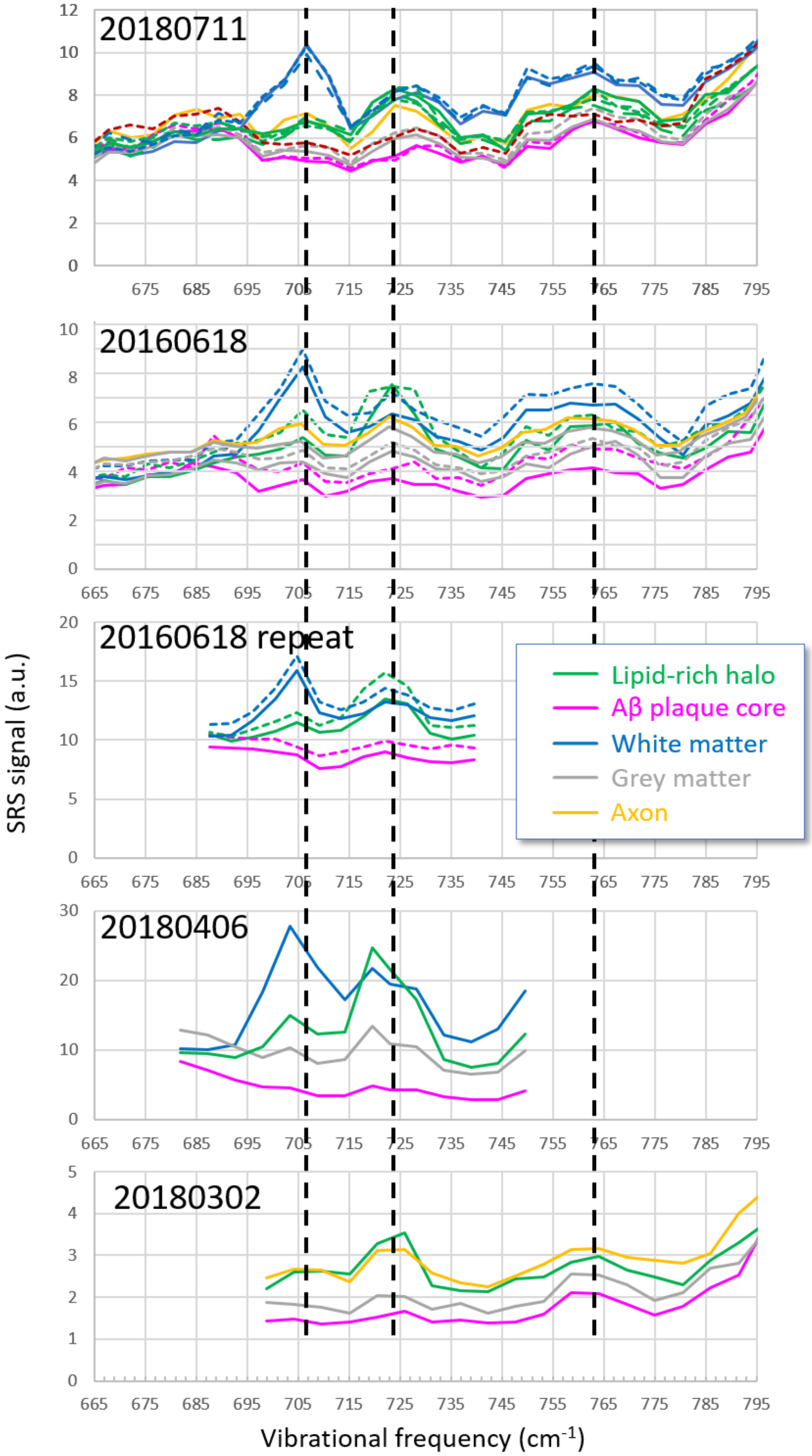


**Supplemental Figure S5:** The robustness and reproducibility of features of signals from Cholesterol, Phosphatidylcholine/Sphingomyelin and Phosphatidylethanolamine is demonstrated by spectra taken on different samples, on multiple different dates.

### Supplemental Figure S6

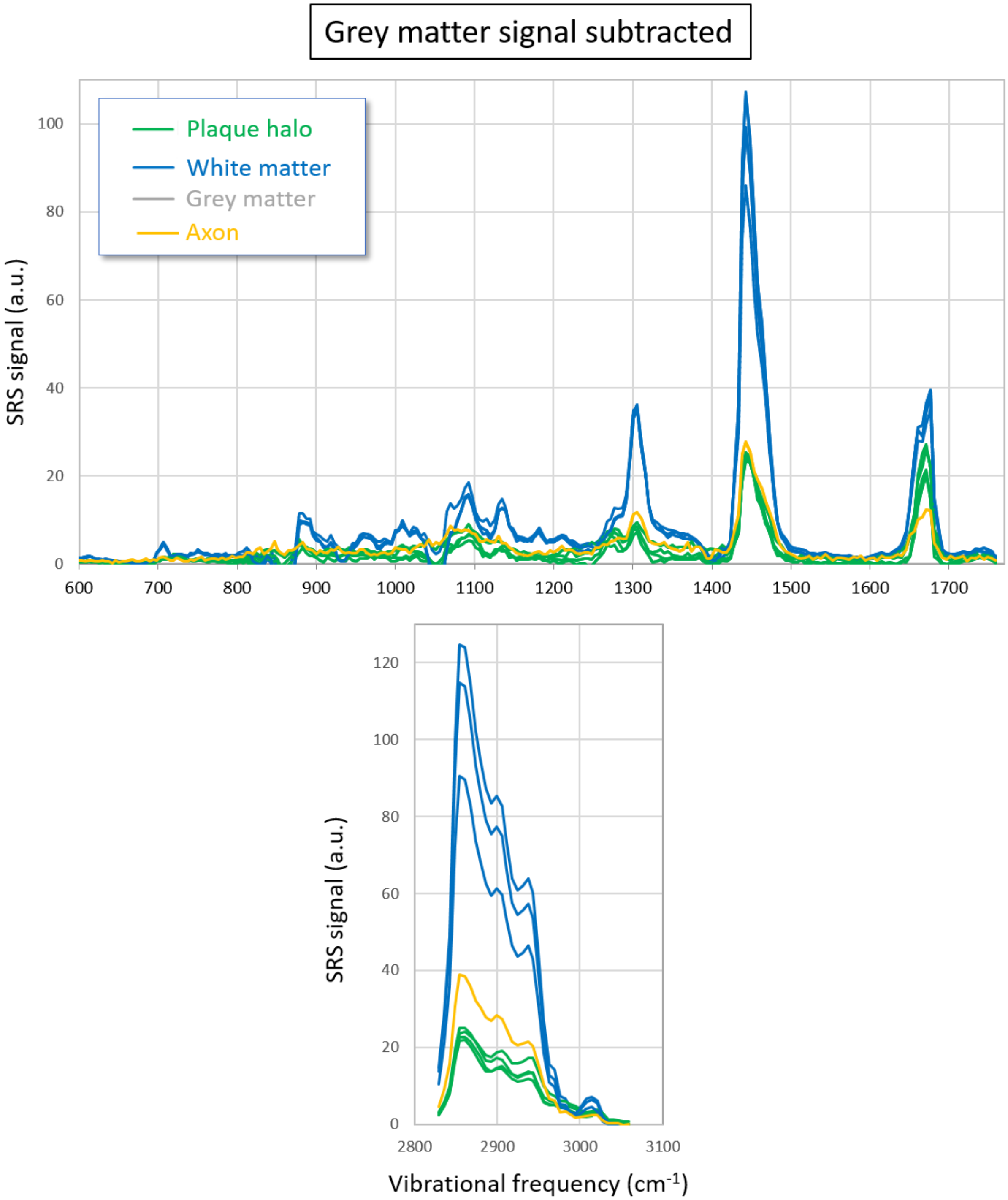


**Supplemental Figure S6:** Grey-matter-subtracted spectra showing preferentially the lipid content of brain structures.

**Supplemental Figure S7**

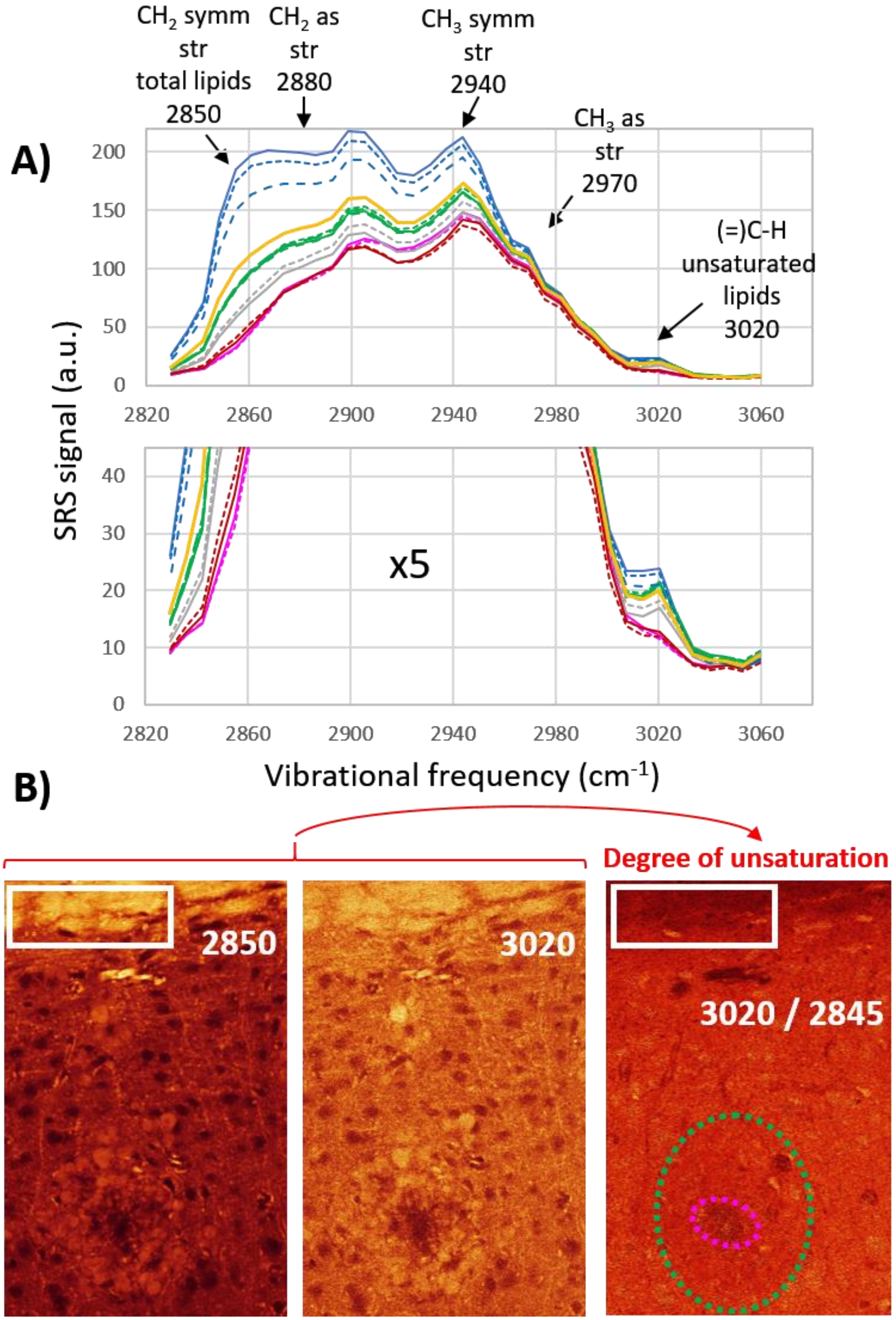


**Supplemental Figure S7:** Information on lipid unsaturation from the high-wavenumber spectral region. (A) High-wavenumber spectra. (B) Ratiometric visualization of unsaturation levels based on =C-H and $CH_2$ stretch signals.
